# Predicting individual incubation of opioid craving by whole-brain functional connectivity

**DOI:** 10.1101/2025.11.08.687395

**Authors:** Ying Duan, Pei-Jung Tsai, Zilu Ma, Ida Fredriksson, Danni Wang, Hanbing Lu, Yavin Shaham, Yihong Yang

## Abstract

A high risk of relapse triggered by craving during abstinence remains a main challenge in opioid addiction treatment. Multiple brain regions have been implicated in opioid craving, but the brain-wide neural mechanisms underlying this process remain poorly understood. Using resting-state fMRI and connectome-based predictive modeling, we identified a whole-brain connectome that predicted the time-dependent increases (incubation) in oxycodone craving in individual rats after voluntary abstinence induced by exposure to an electric barrier. Incubation of oxycodone craving was operationally defined as the increase in non-reinforced lever pressing during relapse tests from early (day 1) to late (day 15) abstinence (incubation score).

We found that changes in whole-brain functional connectivity during abstinence, but not during oxycodone self-administration, predicted the incubation score. Greater decreases in functional connectivity were associated with higher incubation scores. The predictive connectome involved complex interactions across multiple brain systems, including frontal-striatal, frontal-insula, insula-striatal, and hippocampal and sensorimotor circuits. To test causality of the predictive connectome, we examined the effect of pharmacological inactivation of dorsomedial striatum (DMS), which significantly decreased oxycodone seeking after electric barrier-induced abstinence. DMS inactivation increased connectivity strength within the predictive connectome, supporting a causal role of this connectome in incubation of oxycodone craving. The predictive connectome did not predict food-reward seeking after electric barrier-induced abstinence, indicating specificity to oxycodone craving.

Our findings identify a brain-wide connectome marker that predicts individual differences in the incubation of opioid craving and provide potential targets for developing personalized interventions and monitoring therapeutic outcomes in opioid addiction treatment.

**Significance statement:** Relapse driven by craving remains a central challenge in treating opioid addiction. Using resting-state fMRI and connectome-based predictive modeling, we identified a whole-brain connectivity pattern that predicted incubation of oxycodone craving after voluntary abstinence in rats. Connectivity changes during abstinence predicted the magnitude of incubation, measured by increased oxycodone seeking from early to late abstinence. The predictive connectome involved interactions among cortical, basal ganglia, insular, hippocampal and sensorimotor systems. Pharmacological inactivation of the dorsomedial striatum modulated connectivity strength within this network, demonstrating a causal role of the connectome in incubation of craving. These findings identify a brain-wide marker of incubation of opioid craving and provide a translational framework for developing targeted interventions and monitoring therapeutic outcomes in opioid addiction treatment.

## 1. Introduction

Opioid addiction has become an increasingly prevalent and deadly disease in recent decades [1]. Despite the urgent need to address this crisis, available treatment options, such as opioid agonist maintenance therapy, have remained largely unchanged over the past several decades [2]. A major challenge of current treatments is the high risk of relapse, often triggered by craving during abstinence [3].

In humans, abstinence from opioid use is typically voluntary and motivated by the negative consequences of drug seeking [4, 5]. To better model this clinical feature, we recently developed a rat model in which abstinence is induced by adverse consequences of drug seeking—specifically, by requiring rats to cross an electric barrier placed in front of the drug-paired lever [6]. In our initial study using rats trained to self-administer intravenous oxycodone, we found that compared with classical homecage forced abstinence, electric barrier-induced abstinence increases the incubation of oxycodone craving [6]. The term incubation of drug craving refers to the time-dependent increase in non-reinforced drug seeking during abstinence [7, 8].

In subsequent studies, we combined pharmacological, neuroanatomical, and resting-state fMRI methods to identify several brain regions and circuits involved in incubation of oxycodone craving after electric barrier-induced abstinence. These included ventral subiculum, orbitofrontal cortex, claustrum, and dorsomedial striatum (DMS) [9–13]. In our previous resting-state fMRI study, we found that changes in functional connectivity between orbitofrontal cortex and dorsal striatum during abstinence negatively correlate with the strength of incubation of oxycodone craving, quantified by the incubation score—defined as the difference in non-reinforced lever presses between abstinence day 15 (after electric barrier exposure) and day 1 (before barrier exposure) [12]. In a follow-up study, we showed that pharmacological inactivation of the DMS not only decreased relapse to oxycodone seeking after electric barrier-induced abstinence but also restored longitudinal changes in DMS-cortical functional connectivity observed during abstinence [10].

Together, these studies identified several brain regions involved in incubation of oxycodone craving after electric barrier-induced abstinence. However, a limitation of our studies is that we primarily tested region- and projection-specific hypotheses. In both humans and laboratory animals, drug-associated cues and contexts activate multiple interconnected brain regions, inducing drug craving and seeking [14–17]. Additionally, a recent landmark study showed that the magnitude of cue-induced subjective drug craving in humans can be predicted by the whole-brain patterns of neuronal activity elicited by drug-associated cues [16]. These findings underscore the need to identify brain-wide networks that underlie craving to advance mechanistic understanding of drug craving and relapse.

In the present study, we applied a connectome-based predictive modeling (CPM) approach on existing rodent resting-state fMRI datasets [10, 12]. CPM is a data-driven method for identifying brain-behavior relationships [18] and has been used to predict individual differences in sustained attention [19]. More recently, CPM has been applied to human neuroimaging studies to identify neural markers of cocaine and opioid abstinence, showing distinct anatomical substrates for these two drug classes [20, 21]. A recent study using CPM further showed that subjective craving across cocaine and gambling addiction can be predicted from whole-brain connectivity patterns [22].

However, interpretation of brain imaging studies in humans is challenging because participants often receive concurrent treatment interventions, introducing variability and confounds that are difficult to control [21]. Additionally, ethical and practical constraints limit the ability to perform causal manipulations in human studies. In contrast, preclinical fMRI models offer several advantages, including rigorous experimental control, the ability to track longitudinal brain changes during self-administration, abstinence, and relapse, and the opportunity to establish causal relationships through invasive or pharmacological manipulations [23]. These advantages increase mechanistic inference and facilitate translation from preclinical to clinical research.

The goal of our study was to identify a whole-brain functional connectivity pattern (connectome) that would predict the incubation of oxycodone craving. First, we applied CPM to resting-state fMRI data collected during either the oxycodone self-administration phase or the electric barrier-induced voluntary abstinence phase in our rat model [12]. Second, to test for causal role of the identified connectome in incubation, we examined whether pharmacological inactivation of the DMS, previously shown to reduce relapse to oxycodone seeking [10], distinctly modulates the identified connectome relative to the control conditions. Finally, we tested the specificity of the identified connectome by assessing its ability to predict food-reward seeking in a separate group of rats trained to self-administer palatable food and tested for relapse to food seeking after electric barrier-induced abstinence [12].

## 2. Results

### 2.1 Identification of a connectome predictive of incubation of oxycodone craving

We built a connectome-based predictive model [18] to identify the brain connectome that predicts the incubation of oxycodone craving. We analyzed our previously published dataset [12] in which male rats (n = 18) self-administered oxycodone for 14 days and then underwent 12 days of electric barrier-induced abstinence (**Figure 1A**). We collected resting-state fMRI data before oxycodone self-administration training (pretraining) and on abstinence day 2 and 16, one day after the early and late abstinence relapse tests (**Figure 1A**). We used individual changes in functional connectivity during oxycodone self-administration or electric barrier-induced abstinence to predict individual changes in oxycodone craving (incubation score) from early (day 1, before barrier exposure) to late (day 15, after exposure) abstinence. We defined the incubation score as the difference in non-reinforced active lever presses between day 15 and day 1 relapse test.

**Figure 1.**
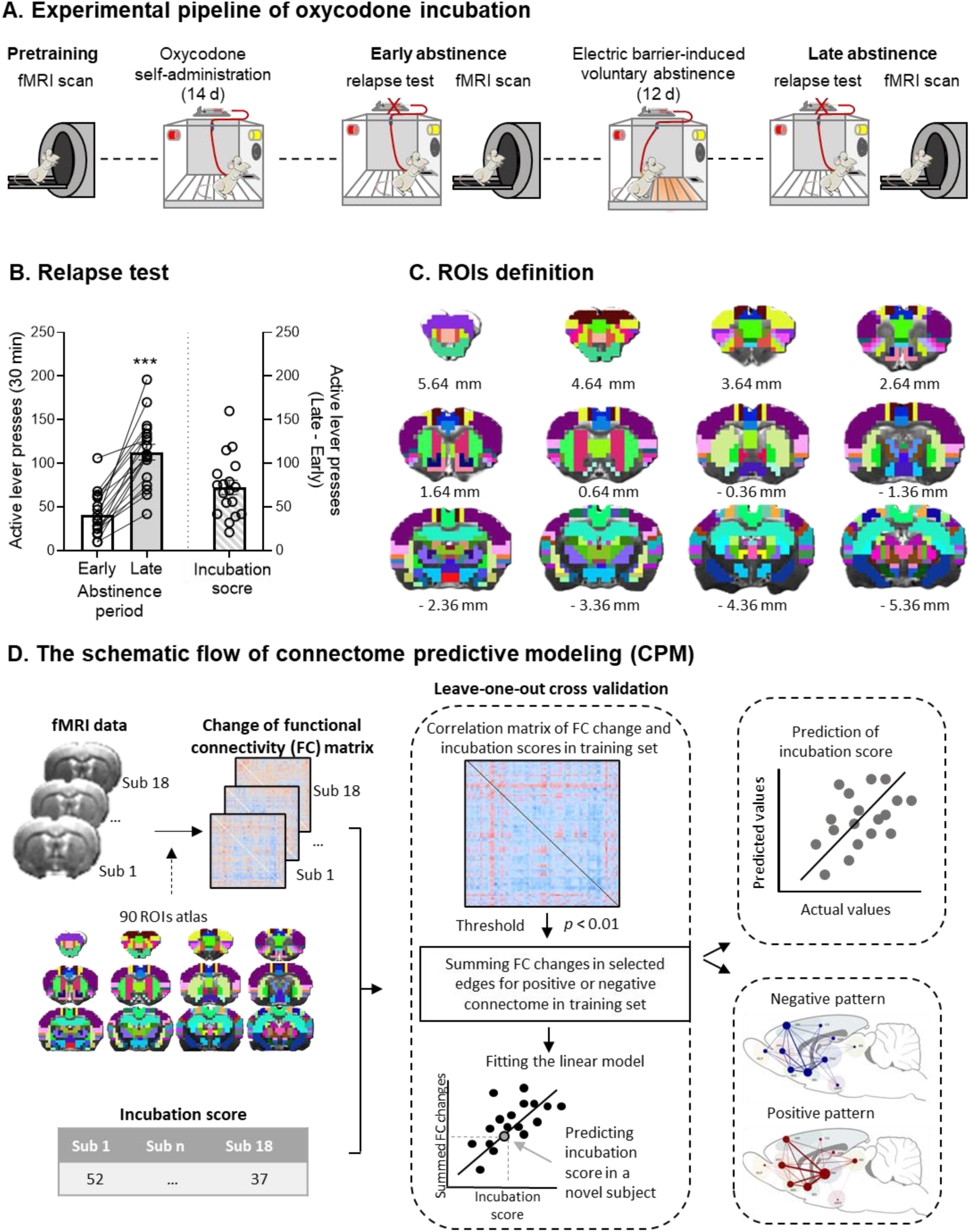
Incubation of oxycodone craving after electric barrier-induced abstinence and connectome-based predictive model framework. (**A**) Timeline of the experiment. (**B**) <u>Relapse (incubation) tests:</u> mean ± SEM and individual data for active lever presses during the 30-minute early abstinence (day 1) test and the first 30 minutes of the late abstinence (day 15) test (left), and mean ± SEM and individual data for the incubation scores, defined as active lever presses during the first 30 minutes of the late abstinence test minus active lever presses during the 30-minute early abstinence test) (right). *** p <0.001, different from Day 1. Data are from ref. [12]. (**C**) The 90 regions of interest (ROIs) according to the Paxinos and Watson [62] rat brain atlas. (**D**) Schematic overview of the connectome-based predictive modeling (CPM) workflow. For each rat, we extracted time series from each ROI and computed pairwise correlations to generate connectivity matrices. We used changes in connectivity matrices across the self-administration and voluntary abstinence phase and the incubation scores as model inputs. Under leave-one-out cross-validation, we held out data from one rat for testing and used the remaining rats for training. Within the training set, we computed Pearson’s r correlations between each edge and the incubation scores. We selected edges with significant correlations (p < 0.01) to form positive and negative connectomes, representing edges with positive or negative correlations with the incubation scores in a novel rat. After 18 iterations, we computed correlations between the predicted and actual incubation score values to assess model performance and identified predictive patterns.

The rats showed reliable incubation of oxycodone craving after electric barrier-induced abstinence [12]. They pressed the active lever significantly more during the 30-minute relapse test on day 15 than on day 1 (paired t-test, t _(17)_ = 8.865, p < 0.001; **Figure 1B**).

We computed whole-brain connectomes (connectivity matrices) for 90 regions-of-interest (ROIs; **Figure 1C**) at pretraining, early abstinence, and late abstinence. We calculated changes in connectivity for the self-administration phase (early abstinence minus pretraining) and the abstinence phase (late minus early abstinence) and used these changes in CPM analyses to predict individual incubation scores (**Figure 1D**).

During voluntary abstinence, the negative connectome (connectivity negatively correlated with the incubation scores; see Methods) significantly predicted incubation scores. **Figure 2A** shows the contribution of each edge to the predictive connectome. Edges involving the frontal cortices, insula, striatum, and sensorimotor cortices showed the strongest weightings. The predicted incubation scores, derived from this predictive connectome, significantly correlated with the actual incubation scores (r = 0.52, p _perm_ = 0.019; **Figure 2B**), indicating successful prediction. In contrast, the positive connectome (connectivity positively correlated with the incubation scores) did not predict incubation (r = −0.31, p _perm_ = 0.16). Changes in connectivity during the self-administration phase also did not predict the incubation score for either the positive (r = 0.15, p _perm_ = 0.34) or negative connectome (r = −0.39, p _perm_ = 0.08). These results showed that only the negative connectome during the abstinence phase predicted the incubation of oxycodone craving.

**Figure 2.**
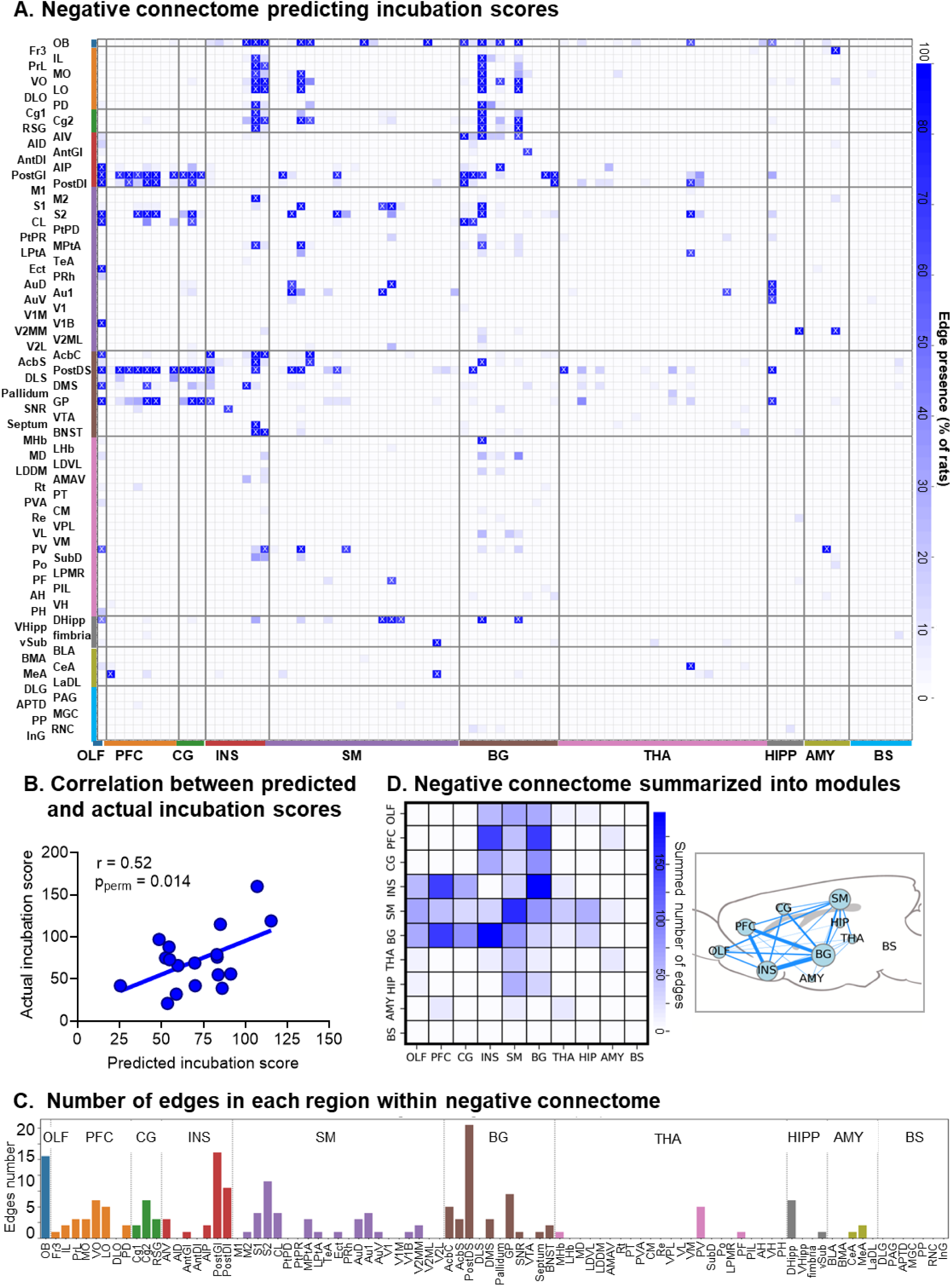
Negative connectome predicting individual incubation of oxycodone craving and model performance. (**A**) The matrix shows edges negatively correlated with the incubation scores (p < 0.01) across all iterations. The color bar indicates the percentage of rats in which each edge appears. Crosses mark edges present in ≥ 50% of rats. We grouped brain regions into 10 anatomical modules: olfactory system (OLF), prefrontal cortex (PFC), cingulate cortex (CG), insula (INS), sensorimotor cortex (SM), basal ganglia (BG), thalamus/hypothalamus (THA), hippocampus (HIPP), amygdala (AMY), and brain stem (BS). Table 1 provides full names of all brain regions. (**B**) Scatter plot shows the correlation between predicted and actual incubation scores (r = 0.52, p _perm_ = 0.013). (**C**) Contribution of each region to the negative connectome by summing edges present in ≥ 50% of rats. (**D**) The negative connectome summarized into 10 anatomical modules reflects surviving edges (present in ≥ 50% of rats) within or between modules (left). The graph visualization places modules in a rat sagittal atlas view (right). Node size reflects the total number of surviving edges and line thickness represents the number of surviving edges between modules.

**Table 1.**
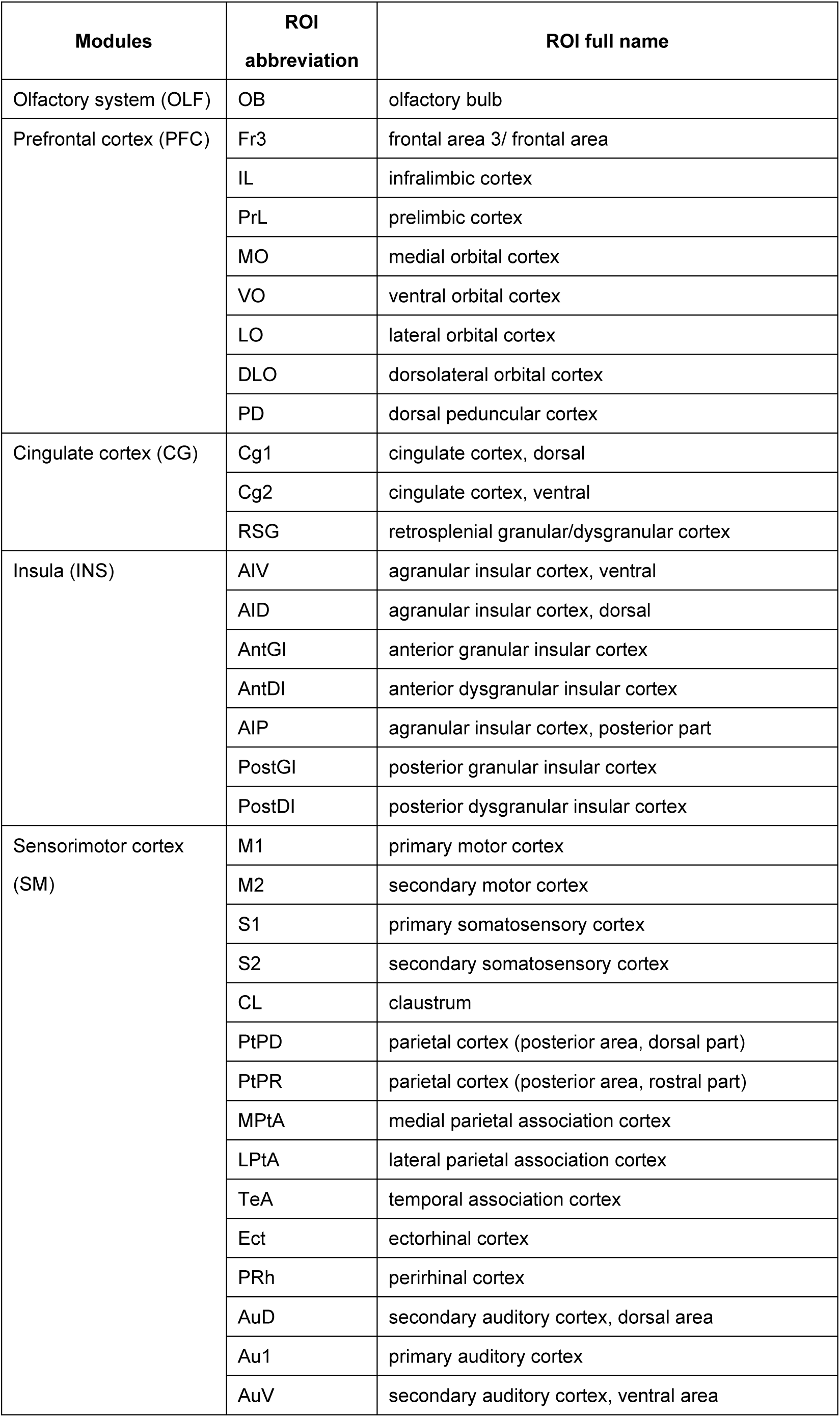
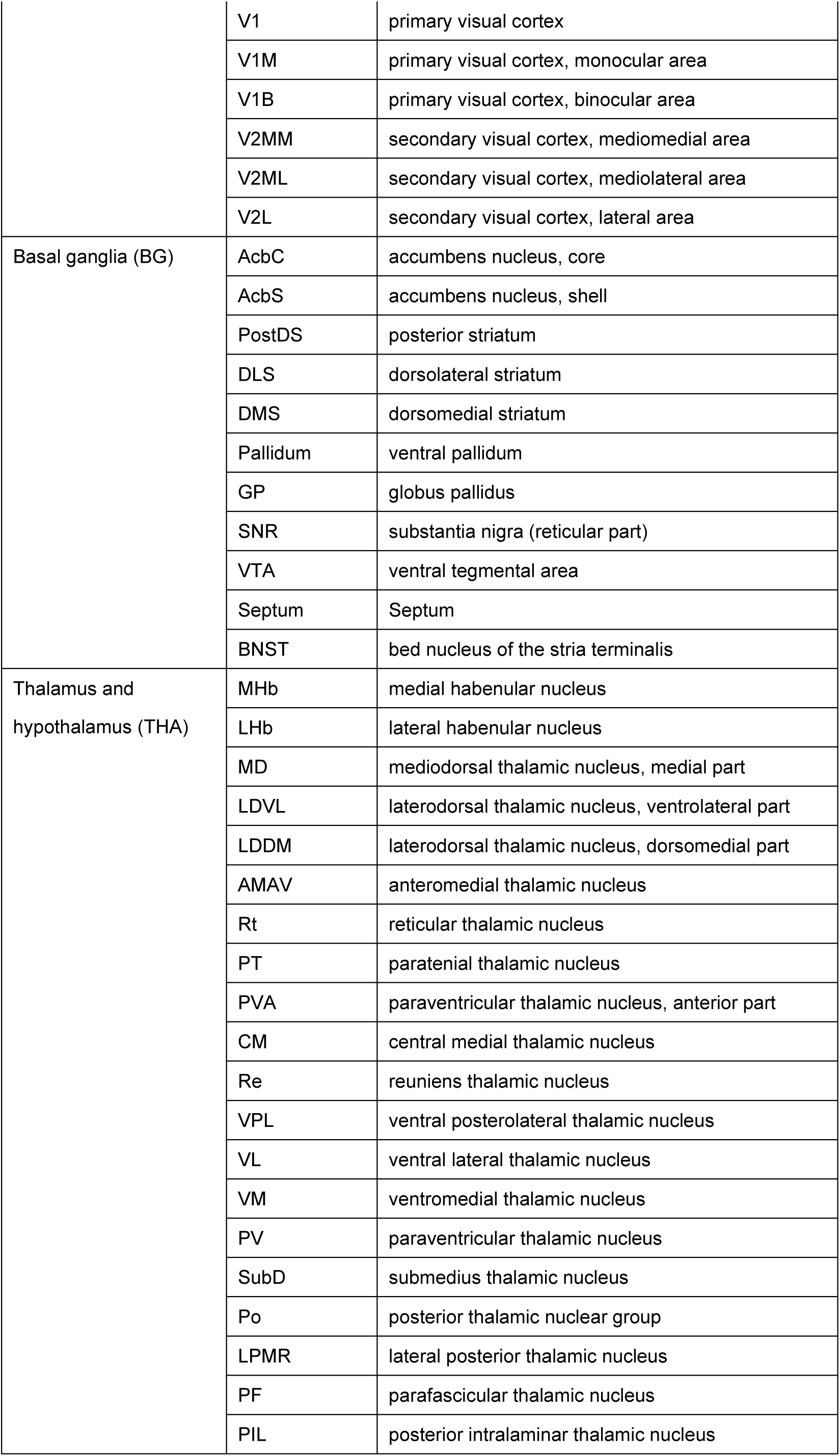
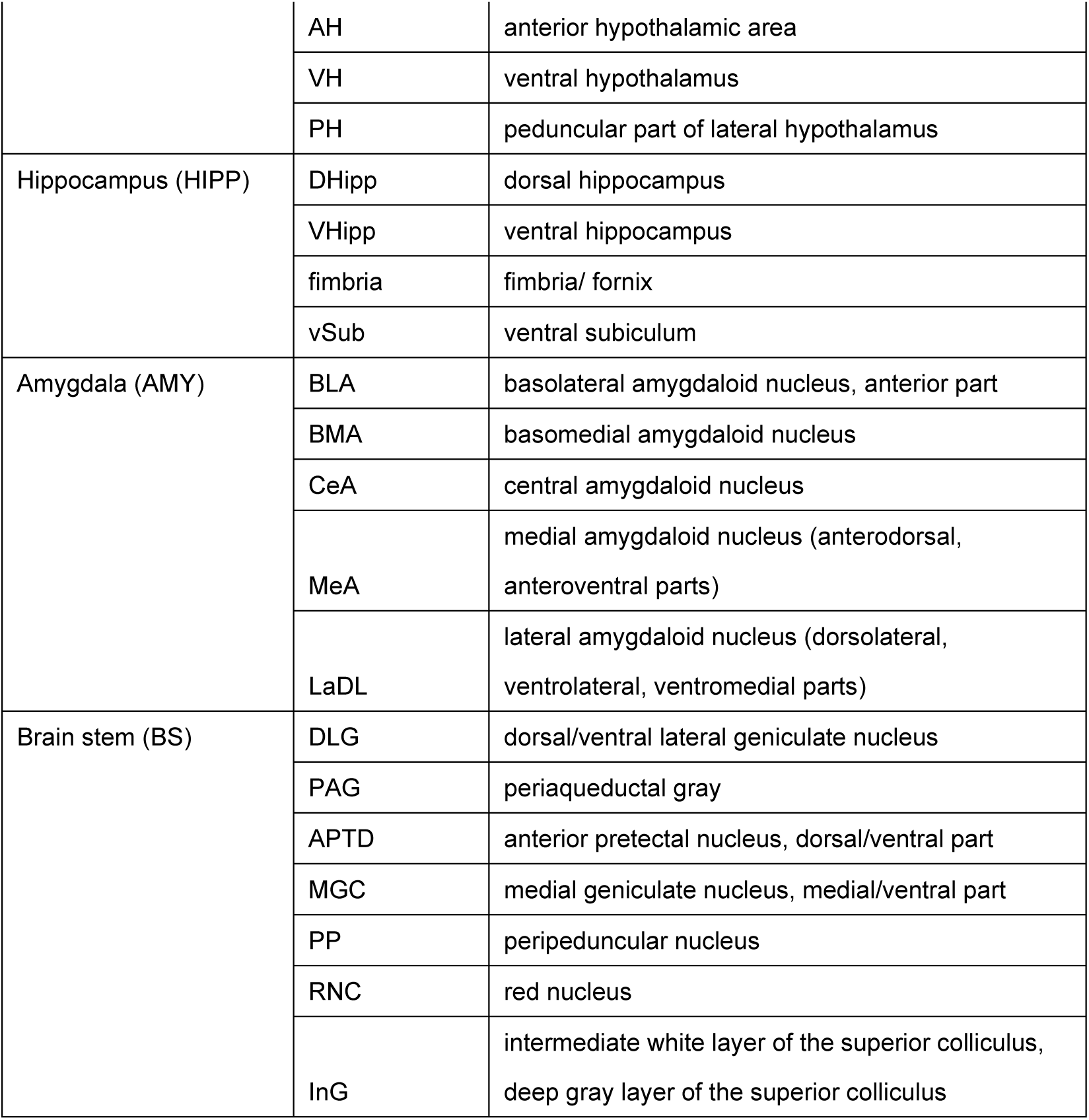
Modules and list of abbreviations.

To identify the brain regions that contributed most to the prediction, we quantified the number of predictive edges connected to each ROI or structural module that appeared in more than 50% of subjects (**Figure 2A**). The top 10 contributing regions included the olfactory bulb, ventral orbitofrontal cortex, cingulate cortex, secondary sensory cortex, posterior granular insula, posterior dysgranular insula, posterior striatum, globus pallidus, and dorsal hippocampus (**Figure 2C**).

At the systems level, connectivity between the prefrontal cortex and insula, between the prefrontal cortex and basal ganglia, between the insula and basal ganglia, and within the sensorimotor module contributed most strongly to the prediction (**Figure 2D**; **Table 1**).

In summary, CPM analysis of whole-brain connectivity changes during voluntary abstinence identified a connectome that predicts individual differences in the incubation of oxycodone craving. Changes in connectivity among prefrontal, insular, basal ganglia, and sensorimotor regions primarily drove this prediction

### 2.2 Causal validation of connectome via intracranial regional inactivation

We next tested whether the identified connectome reflected core features of incubation of craving by analyzing a prior dataset that examined the effect of the DMS inactivation on oxycodone seeking after electric barrier induced abstinence [10]. The experimental design matched that of the connectome identification study [12], except that one group received an intracranial injection of muscimol + baclofen (GABA receptor agonists; n = 9) or saline (n = 12) into the DMS before the late abstinence relapse test. Another group (n = 6 per condition, male) received the same injections after electric barrier-induced abstinence before fMRI scanning (**Figure 3A**).

**Figure 3.**
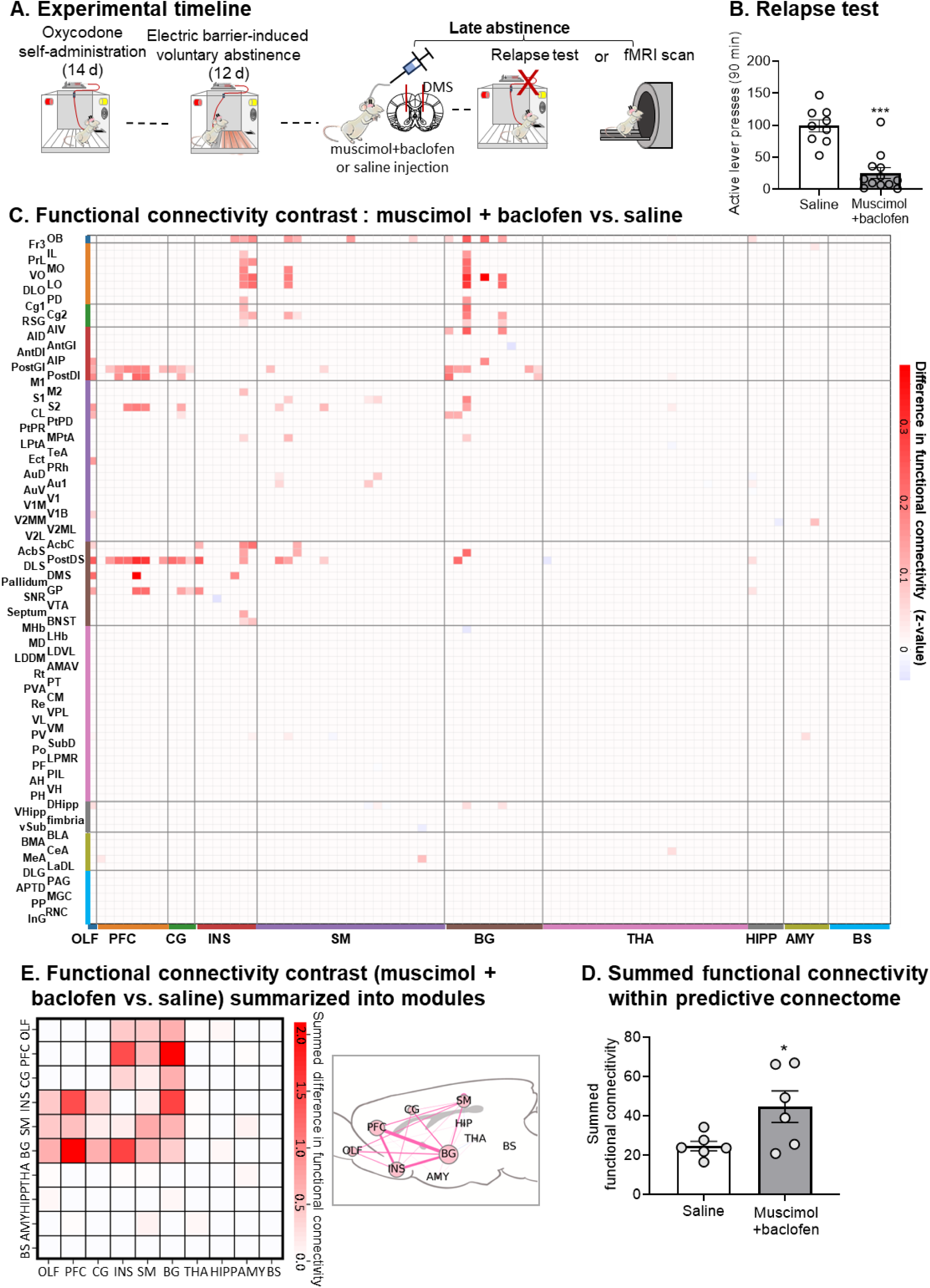
Effect of dorsomedial striatum (DMS) inactivation on the predictive connectome after electric barrier-induced voluntary abstinence. (**A**) Timeline of the experiment. (**B**) <u>Relapse (incubation) test day 15:</u> mean ± SEM and individual data for active lever presses during the 30-minute test after pretreatment with saline (n = 9) or muscimol + baclofen injections into the DMS (n = 12). *** p < 0.001, different from saline. Data are from ref. [10]. (**C**) Heatmap illustrates the contrast in functional connectivity between the muscimol + baclofen and saline groups (muscimol + baclofen minus saline) across edges within the predictive connectome. The bottom annotation shows the 10 modules indicating clustered regions. (**D**) The heatmap shows the difference in functional connectivity within the predictive connectome between muscimol + baclofen and saline injections summarized into 10 modules (left). A graph visualization places modules in a rat sagittal atlas view (right). Node size represents the summed difference in functional connectivity (muscimol + baclofen minus saline) within and between modules, and edge thickness represents the summed difference in functional connectivity between modules. These results showed that DMS muscimol + baclofen injections increased connectivity within and between modules within predictive connectome. (**E**) Bar plot shows summed functional connectivity within the predictive connectome after saline or muscimol + baclofen injections into the DMS. Muscimol + baclofen injections significantly increased functional connectivity compared with saline (* p < 0.05).

Muscimol + baclofen inactivation of the DMS significantly reduced oxycodone seeking during late abstinence [10] (unpaired t-test, t _(19)_ = 5.77, p < 0.001; **Figure 3B**). We then compared functional connectivity within the predictive connectome (edges present in ≥50% of subjects) between treatment groups. The muscimol + baclofen group exhibited higher overall functional connectivity (**Figure 3C**). This group showed higher connectivity between the prefrontal cortex and insula, between the prefrontal cortex and basal ganglia, and between the insula and basal ganglia modules compared to the saline group (**Figure 3D**). Additionally, the muscimol+ baclofen group showed significantly greater summed connectivity strength within the predictive connectome (unpaired t-test, t_(10)_ = 2.38, p = 0.039; **Figure 3E**).

Together, these results show that muscimol + baclofen inactivation, which inhibits incubated oxycodone seeking [10], increased functional connectivity within the predictive connectome. Higher connectivity corresponded to lower incubated oxycodone seeking after electric barrier-induced abstinence. This pattern of results indicates that the functional connectome plays a causal role in incubation of oxycodone craving.

### 2.3 Specificity of connectome predictive of oxycodone but not food craving

We tested the specificity of the predictive connectome by applying it to a food control group (n = 19, males) that underwent parallel procedures to those of the oxycodone-experience rats [12], including food self-administration, electric barrier-induced abstinence, early and late relapse tests, and fMRI scanning (**Figure 4A**).

**Figure 4.**
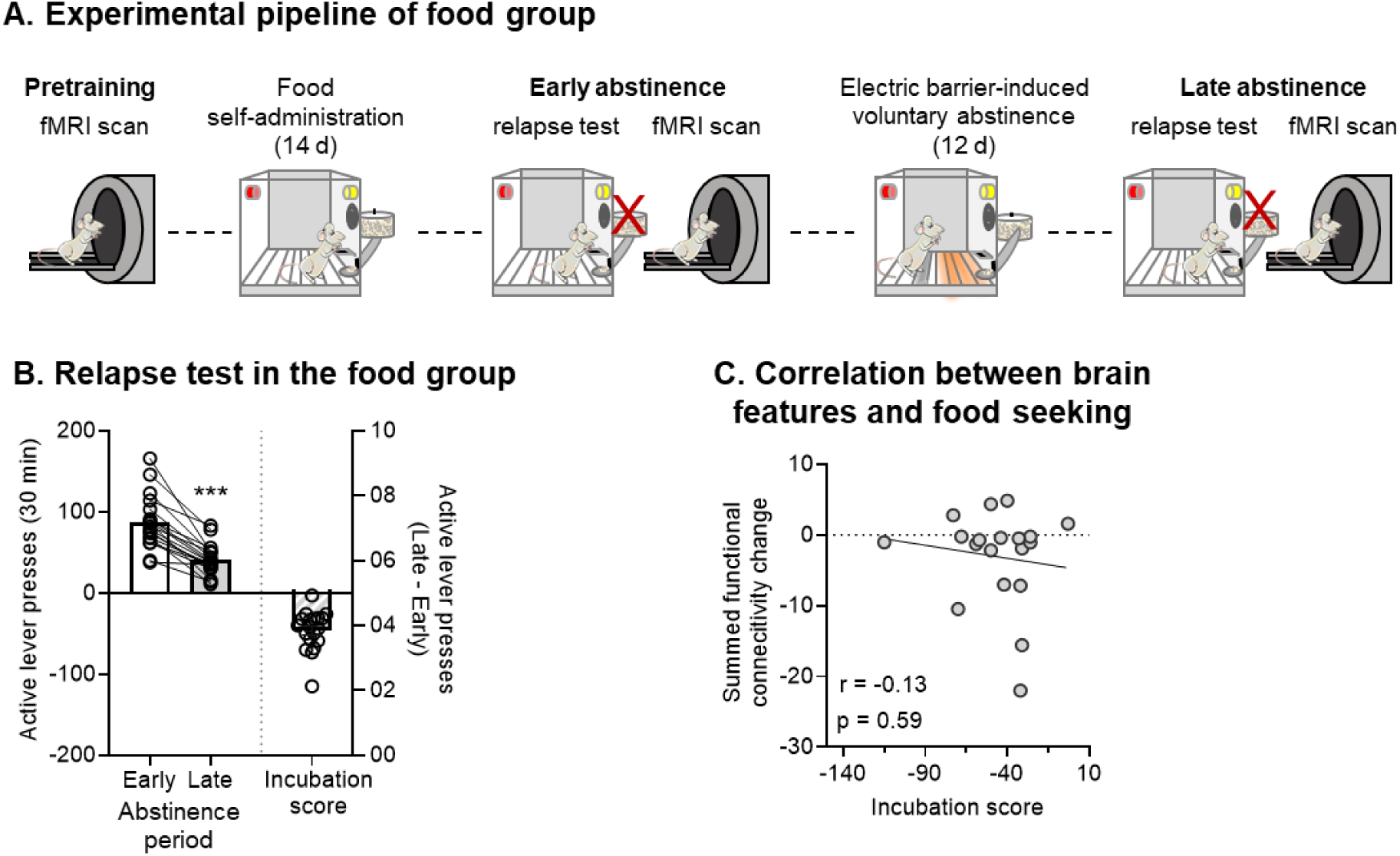
Performance of the identified predictive connectome in drug-naïve rats with a history of food self-administration. (**A**) Timeline of the experiment. (**B**) <u>Relapse (incubation) tests:</u> mean ± SEM and individual data for active lever presses during the 30-minute early abstinence (day 1) test and the first 30 minutes of the late abstinence (day 15) test (left), and mean ± SEM and individual data for the incubation score (active lever presses during the first 30 minutes of the late abstinence test minus active lever presses during the 30-minute early abstinence test) (right). *** p <0.001, different from Day 1. Data are from ref. [12]. (**C**) Scatter plot shows the relationship between summed functional connectivity changes in the predictive connectome and the food incubation scores after electric barrier-induced abstinence. We found no significant correlation between the connectome and the food incubation score (r = −0.13, p = 0.59).

Unlike the oxycodone group, the food group decreased food seeking from early to late abstinence (paired t-test, t_(20)_ = 8.34, p < 0.001; **Figure 4B**). We summed the change in functional connectivity during abstinence within the predictive connectome (edges observed in ≥50% of rats; **Figure 2A**) and correlated this measure with individual changes in food seeking (late minus early abstinence, the incubation score). This analysis showed no significant relationship (r = −0.13, p = 0.59; **Figure 4C**), indicating that the predictive connectome does not predict time-dependent changes in food craving and seeking.

Together, the results indicate that the identified connectome serves as a specific neural marker for opioid-related craving and relapse, distinct from neural circuits underlying craving for and seeking of food rewards.

## 3. Discussion

We identified a predictive connectome for incubation of oxycodone craving in a rat model. Our study has three main findings. First, individual changes in whole-brain functional connectivity during the abstinence phase, but not the self-administration phase, predicted individual incubation scores. The identified connectome negatively correlated with incubation and included large-scale neural networks, including the prefrontal cortex, insula, basal ganglia, and sensorimotor structures. Second, in an independent dataset, pharmacological inactivation of the DMS, previously shown to decrease incubation of oxycodone seeking after electric barrier-induced abstinence [10], increased functional connectivity within the predictive connectome. This result suggests a causal role of the predictive connectome in incubation of opioid craving. Third, applying the same predictive features to a food control group did not predict food seeking after electric barrier-induced abstinence, indicating that the connectome is specific to oxycodone craving.

### 3.1 Functional connectivity changes during voluntary abstinence predict incubation of craving

Using a data-driven CPM approach, we identified a negative predictive connectome during the voluntary abstinence, but not during the self-administration phase. This finding agrees with our previous seed-based, hypothesis-driven analyses showing that changes in orbitofrontal cortex-striatal and claustrum-cortical functional connectivity during voluntary abstinence phase, but not the self-administration phase, negatively correlate with the incubation score [11, 12]. These results highlight the temporal sensitivity of functional connectivity changes, suggesting that craving-related neural adaptations emerge specifically during abstinence rather than during opioid exposure.

This observation is consistent with molecular and behavior evidence showing unique neuroadaptations during periods of forced or voluntary abstinence that are critical to incubation of drug craving [7, 24]. For example, long-term forced abstinence recruits calcium-permeable AMPA receptors (CP-AMPARs) receptors in nucleus accumbens, and pharmacological blockade of CP-AMPARs selectively decreases incubated cocaine or oxycodone seeking after prolonged abstinence [25, 26].

Together, these findings suggest that abstinence represents a critical window for the formation of craving-related brain reorganization. Determining the causal relationship between cellular adaptations and large-scale functional connectivity changes is a subject for future research.

### 3.2 Key neural circuits within predictive connectome that contribute to incubation

The predictive connectome includes functional connectivity among frontal cortices, basal ganglia, and sensorimotor cortices, highlighting complex interactions that support the neural mechanisms underlying incubation of opioid craving. Theoretical frameworks of addiction emphasize drug-induced changes in cortico-striato-pallido-thalamo-cortical loops [27–30]. Our predictive connectome contains multiple cortical-basal ganglia circuits, consistent with these frameworks.

Changes in functional connectivity between orbitofrontal cortex and DMS contributed to the predictive connectome, in agreement with our previous results [10, 12]. These regions are critical for incubation of heroin and oxycodone craving in rats, as pharmacological inactivation of either region decreases incubation [10, 31, 32]. Supporting these findings, exposure to opioid-associated cues increases fMRI signal in humans with opioid addiction [33, 34].

Beyond prefrontal-striatal circuits, posterior insula-striatal circuits also contributed to the predictive connectome. This observation agrees with a human study showing that decreased connectivity between posterior insula and putamen predicts relapse to cocaine use [35]. The posterior insula integrates interoceptive signals [36] and has been linked drug craving [37]. In rats, posterior insula inactivation decreases incubation of heroin craving [38], suggesting that insula-striatal circuits mediate craving via interoceptive signal processing. Prefrontal-insula connectivity to the predictive connectome further suggest convergence of cognitive and interoceptive processes [39] that may be involved in incubation of opioid craving.

The predictive connectome also includes the central amygdala, a region critical for incubation of drug craving after homecage forced abstinence, and for the protective effect of rewarding social interaction on incubation [40–43]. Connectivity between central amygdala and paraventricular thalamus contributed to the predictive connectome; the latter region has been implicated in relapse to heroin seeking in different rodent models [44–46]. Connectivity between the bed nucleus of stria terminalis (BNST) and insula also contributed to the predictive connectome. The BNST is critical for negative affective states during opioid withdrawal [47] and stress-induced reinstatement of heroin seeking [48, 49]. However, the relationship between these behaviors and incubation remains unclear, particularly as opioid withdrawal and incubation follow opposing time courses [50].

We also identified sensorimotor cortices and hippocampus in the predictive connectome. A human study reported that relapse to cocaine use is associated with increased cue-induced activation of sensory association and motor cortical areas [51]. A recent study in rats, implicates the supplementary motor cortex in incubation of cocaine craving after homecage forced abstinence [52]. Whole brain Fos mapping of incubation of palatable food seeking after homecage forced abstinence also shows that incubation is associated with somatosensory and motor cortices activation [53]. Although the hippocampus has not been directly linked to incubation, it plays a role in context-induced reinstatement across drug classes [54–56].

Finally, the claustrum, which is critical to incubation of oxycodone after electric barrier-induced abstinence [11], is also part of the predictive connectome. Functional connectivity between claustrum with olfactory bulb and cingulate cortex carried significant weighting, partially overlapping with our prior claustrum seed-based analyses [11]. However, connectivity between ventral subiculum (vSub) and retrosplenial cortex, previously found to be associated with incubation [9], did not appear in the current connectome, likely due to differences in ROI definition of vSub: bilateral in the current study vs. unilateral seed in the previous study.

### 3.3 Validation of the predictive connectome and treatment responsiveness

We validated the predictive connectome using an independent group of rats in which we pharmacologically inactivated the DMS. This manipulation decreased oxycodone seeking after electric barrier-induced abstinence [10] and increased DMS functional connectivity with cortical, insular, and basal ganglia regions. These results support a causal relationship between connectome integrity and incubation of oxycodone craving and show that the identified connectome responds to intervention. This finding suggests that the connectome may serve as a functional neural marker (see ref. [16] to assess treatment efficacy, particularly in clinical contexts where subjective craving reports are often not reliable [57].

Finally, applying the connectome to drug-naïve rats with a history of food self-administration showed no significant relationship between functional connectivity and the incubation score, indicating that the identified connectome selectively predicts opioid-related craving and seeking. These results support the translational potential of the connectome for identifying drug-specific treatment targets and monitoring addiction-related circuitry over time.

### 3.4 Conclusion

Using whole-brain resting-state fMRI and a CPM framework, we predicted incubation of oxycodone craving based on functional connectivity changes during voluntary abstinence. The identified connectome encompasses multiple brain regions and circuits, including previously unrecognized contributors such as the BNST–insula and central amygdala–paraventricular thalamus connections. Pharmacological inactivation provided causal evidence, highlighting the connectome’s potential as a target for interventions and for monitoring treatment effects. The connectome’s specificity for oxycodone craving, but not food craving, further supports translational relevance. Overall, we identified a brain-wide neural marker predictive of individual incubation of oxycodone craving, with potential applications in guiding personalized interventions and monitoring therapeutic outcomes in opioid addiction.

## 4. Methods and Materials

### 4.1 Animals

We used 70 Sprague-Dawley rats (Charles River or ENVIGO) weighing 260–360 g before surgery. Before surgery, we housed rats two per cage and then individually afterward under a reverse 12:12-hour light/dark cycle, with food and water freely available. All experimental procedures followed the NIH Guide for the Care and Use of Laboratory Animals and an approved protocol from the Animal Care and Use Committee of our institute.

### 4.2 Experimental Overview

We conducted three sets of analyses on existing datasets [10, 12] to identify and validate the connectome that predicts incubation of oxycodone craving after electric barrier–induced abstinence.

#### Analysis 1: Identification of connectome predictive of incubation of oxycodone craving

We used data from our previously published study [12]. Eighteen male rats underwent 14 days of oxycodone self-administration, 12 days of electric barrier-induced abstinence (voluntary abstinence), and relapse tests on early and late abstinence (days 1 and 15). We collected resting-state fMRI data before self-administration training and on the days after relapse tests (**Figure 1A**). These data allowed us to calculate changes in functional connectivity during both the self-administration and the voluntary abstinence phases. We then used CPM to identify the pattern of functional connectivity change (or connectome change) during these phases that predicted the incubation score (non-reinforced lever presses during the late abstinence relapse test minus lever presses during the early abstinence test).

#### Analysis 2: Causal validation of connectome via intracranial regional inactivation

We tested whether pharmacological inactivation of the DMS, which suppressed incubation of oxycodone craving after voluntary abstinence [10], affected the predictive connectome identified in Analysis 1. We used a dataset that examined the effect of DMS inactivation on incubation of oxycodone craving [10]. Rats underwent 14 days of oxycodone self-administration and 14 days of electric barrier–induced abstinence. On abstinence day 15, we injected rats with muscimol + baclofen or saline 30 min before either relapse testing (behavioral cohort, n = 21, male and female) or fMRI (imaging cohort, n = 12, male) (**Figure 3A**).

#### Analysis 3: Specificity of connectome predictive to oxycodone but not food craving

To evaluate whether the identified predictive connectome specifically predicted oxycodone craving, we analyzed another dataset in which 19 male rats self-administered food instead of oxycodone as a control group [12]. We followed the same procedures used in Analysis 1 (**Figure 4A**) to test whether the oxycodone-predictive connectome generalized to the prediction of food craving. Below, we describe the general procedures for self-administration, voluntary abstinence, relapse testing, and fMRI acquisition and preprocessing. We then describe experiment-specific analyses in separate subsections. For additional details of the surgical and behavioral procedures, see ref. [9, 10].

### 4.3 General Procedures

#### Intravenous surgery

We anesthetized rats with isoflurane (5% induction, 1.5–2.5% maintenance) and inserted Silastic catheters attached to a modified 22-gauge cannula on polypropylene mesh into the jugular vein. We secured the catheter and administered ketoprofen (2.5 mg/kg) after surgery and again the next day for analgesia. Rats recovered for 6–8 days before self-administration training. We flushed catheters every 24–48 hours with gentamicin and heparin [10, 12].

#### Intracranial surgery and injection

For the dataset in Analysis 2, after intravenous catheterization, we placed the rats in a stereotaxic frame and implanted bilateral guide cannulas 1.5 mm above the dorsomedial striatum (AP +0.6, ML ±2.1, DV –4.2, relative to bregma). We secured the cannulas with dental cement. We diluted muscimol and baclofen (Sigma-Aldrich, M1523 and B5399) in sterile saline and injected 50 + 50 ng in 0.5 µl into the dorsomedial striatum 30 minutes before the relapse test (behavioral cohort) or fMRI scanning (imaging cohort) [10]. The doses of muscimol + baclofen are based on our previous studies [9, 58, 59].

#### Self-administration training

Rats self-administered oxycodone (0.1 mg/kg/infusion) or 45 mg palatable food pellets [60] for 6 hours per day (six 1-h sessions separated by 10 min) for 14 days. Each session began with houselight illumination, followed by active lever insertion 10 seconds later. Active lever presses delivered drug or food rewards paired with a 20-second tone-light cue under a fixed-ratio 1 schedule with a 20-second timeout. Inactive lever presses were recorded but had no consequence. Sessions ended with lever retraction and lights off. We limited the number of oxycodone infusions or food pellet rewards to 15 per hour.

#### Electric barrier–induced voluntary abstinence

During the 12-day electric barrier-induced abstinence phase, we made rewards (oxycodone or food) available for 2 hours per day under the same parameters used during training. We induced abstinence by placing an electric barrier (“shock zone”) in front of the reward lever, separated from a “safe zone” by a plastic divider. The shock zone delivered a continuous mild foot shock (0.1–0.4 mA). Current began at 0 mA on day 1 and increased by 0.1 mA per day to 0.3 mA, with an increase to 0.4 mA if intake remained greater than three rewards per day.

#### Relapse tests

On abstinence days 1 or 15, we tested the rats for relapse to oxycodone or food seeking under extinction conditions. After a 30-minute habituation period in the self-administration chamber, the rats underwent a 30-minute (day 1) or 90-minute (day 15) relapse test. Lever presses produced the reward-paired tone-light cue, but no drug or food.

#### MR image acquisition and preprocessing

We acquired high-resolution T2-weighted anatomical images and resting-state fMRI scans using a Bruker Biospin 9.4T MRI scanner. Throughout scanning, we anesthetized rats with dexmedetomidine and isoflurane, positioned them in a prone position, secured them with customized equipment, and continuously monitored physiological parameters. We reported detailed acquisition parameters in ref. [10, 12]. We processed all functional images with a standardized pipeline that included skull stripping, motion correction, coregistration to a template, noise component removal, band-pass filtering (0.01– 0.1 Hz), and spatial smoothing (full width at half maximum = 0.6 mm) [10, 12, 61].

#### Functional connectivity matrix

We parcellated the whole brain into 90 regions of interest (ROIs) using the Paxinos and Watson rat atlas (**Figure 1C**) [62]. From preprocessed resting-state data, we obtained time courses for each ROI and calculated Pearson’s correlation (r values) between each pair of ROIs to generate functional connectivity matrices. We normalized the correlations using Fisher’s z-transformation. We calculated changes in functional connectivity during the self-administration phase (subtracting early abstinence from pretraining) and the voluntary abstinence phase (subtracting late abstinence from early abstinence). This process produced two 90 × 90 matrices of connectivity changes per rat, with each element (edge) representing the change in connectivity between ROIs during the different phases. To characterize the identified networks, we categorized the 90 ROIs into 10 anatomical modules: olfactory system, prefrontal cortex, cingulate cortex, insula, basal ganglia, hippocampus, amygdala, sensorimotor cortex, thalamus and hypothalamus, and brain stem (see **Table 1** for details).

### 4.4 Experiment-Specific Analyses

#### Analysis 1: Identifying connectome predictive of incubation of oxycodone craving using connectome-based predictive modeling (CPM)

We built a CPM [18] to identify the connectome that predicts oxycodone craving. We used individual changes in whole-brain connectivity during the self-administration and voluntary abstinence phases and individual changes in craving from early to late abstinence (incubation scores). We calculated connectivity change matrices for each rat during the self-administration phase (early abstinence minus pretraining) and the voluntary abstinence phase (late abstinence minus early abstinence). We measured incubation score by number of active lever presses during the 30-minute relapse test at late abstinence minus those at early abstinence (**Figure 1B**) to reflect the time-dependent increase of opioid seeking after electric barrier–induced voluntary abstinence [9, 12].

As shown in **Figure 1D**, the CPM analysis used a leave-one-out cross-validation approach. We trained the model on n–1 subjects’ connectivity-change matrices and incubation scores and tested it on the left-out subject. In the training dataset, we correlated the connectivity changes in each pair of ROIs (edges) with the incubation scores across subjects. We retained edges significantly correlated with incubation scores (p < 0.01) as predictive features and separated them into positive and negative connectomes. We obtained single-subject aggregate connectivity by summing the weights of positively or negatively correlated edges in the respective connectome. We then fitted linear models between these aggregate connectivity values and the incubation scores in the training set and tested them on the left-out rat.

After leave-one-out cross-validation, we used a permutation test to evaluate the statistical significance of the correlation between predicted and observed incubation scores [18, 19]. To break the true brain-behavior relationship, we randomly shuffled the actual incubation scores across subjects. For each permutation, we applied the same procedure for model training and prediction, and computed the correlation (r) between the predicted and permuted incubation scores. This process was repeated 1,000 times to generate a null distribution of r-values for both the positive and negative connectome. We then compared the observer Pearson correlation from the actual data to the null distribution to calculate the empirical p-value.

To further validate and assess the specificity of the predictive features, we built a predictive connectome that retained only edges consistently observed in more than 50% of subjects (9 out of 18 rats). This consensus-based approach decreased subject-specific variability and ensured that the resulting connectome captured stable and reproducible connectivity most relevant to prediction.

#### Analysis 2: Causal validation of connectome via pharmacological inactivation of DMS

To validate the predictive connectome identified in Analysis 1, we tested the effect of DMS inactivation in a behavioral and an imaging cohort. In the behavioral cohort, we examined whether muscimol + baclofen inactivation of the DMS affected oxycodone seeking after electric barrier– induced abstinence by comparing active lever presses during the 90-minute relapse test between saline and muscimol + baclofen groups using a two-sample t-test as described in ref. [10].

In the imaging cohort, we determined whether pharmacological inactivation modulated the predictive features of the connectome identified in Analysis 1. From preprocessed fMRI data, we constructed connectivity matrices across the same 90 ROIs following either saline or muscimol + baclofen injection. We then calculated the sum of functional connectivity values across edges in the predictive connectome derived from Analysis 1 and compared these sums between the saline and muscimol + baclofen groups using a two-sample t-test.

#### Analysis 3: Specificity of identified connectome predictive of oxycodone but not food seeking

We further evaluated the specificity of the predictive connectome by applying its predictive features to a food control group that underwent the same experimental procedures, except that 45 mg food pellets replaced oxycodone as the operant reward [12]. In the behavioral assessment, we compared active lever presses during the 30-minute relapse test between early and late abstinence using a paired t-test to evaluate food seeking after voluntary abstinence [12].

To test whether the predictive connectome identified in Analysis 1 predicted food seeking, we constructed functional connectivity matrices with the same 90 ROIs. We then obtained connectivity change matrices by subtracting connectivity during early abstinence from that during late abstinence. After summing connectivity changes within the predictive connectome mask, we assessed the correlation between summed functional connectivity changes (late minus early abstinence) and the food incubation score (late minus early abstinence).

## Data and materials availability

The behavioral and imaging data are available upon request (Y.Y.).

